# Thermal decomposition of the amino acids glycine, cysteine, aspartic acid, asparagine, glutamic acid, glutamine, arginine and histidine

**DOI:** 10.1101/119123

**Authors:** Ingrid M. Weiss, Christina Muth, Robert Drumm, Helmut O.K. Kirchner

## Abstract

Calorimetry, thermogravimetry and mass spectrometry were used to follow the thermal decomposition of the eight amino acids G, C, D, N, E, Q, R and H between 185°C and 280°C. Endothermic heats of decomposition between 72 and 151 kJ/mol are needed to form 12 to 70 *%* volatile products. This process is neither melting nor sublimation. With exception of cysteine they emit mainly H_2_O, some NH3 and no CO2. Cysteine produces CO2 and little else. The reactions are described by polynomials, AA ^ a (NH_3_) + b (H_2_O) + c (CO2) + d (H_2_S) + e (residue), with integer or half integer coefficients. The solid monomolecular residues are rich in peptide bonds.

## 1. Motivation

Amino acids might have been synthesized under prebiological conditions on earth or deposited on earth from interstellar space, where they have been found [Follmann and Brownson, 2009]. Robustness of amino acids against extreme conditions is required for early occurrence, but little is known about their nonbiological thermal destruction. There is hope that one might learn something about the molecules needed in synthesis from the products found in decomposition. Our experimental approach is not biochemical, it is merely thermochemical.

## 2. Experimental procedures

### 2.1 DSC, TGA, QMS

Altogether 200 samples of amino acids of at least 99.99 % purity from Sigma-Aldrich were tested in a Simultaneous thermal analysis apparatus STA 449 Jupiter (Netzsch, Selb, Germany) coupled with Mass spectrometer QMS 403C Ae ëolos (Netzsch). Specimens of typically 10 mg weight in Al cans were evacuated and then heated at 5K/min in argon flow. Differential scanning calorimetry (DSC) and thermal gravimetric analysis / thermogravimetry (TGA, TG), as well as quantitative mass spectrometry (QMS) outputs were smoothed to obtain the data of section 3.1. The mass spectrometer scanned 290 times between 30°C and 320°C, i.e. at every single degree in 1 Da steps between 1 Da and 100 Da. Alltogether, 290 x 100 x 200 = 5.8 million data points were analyzed.

### 2.2 Visuals

A MPA120 EZ-Melt Automated Melting Point Apparatus (Stanford Research Systems, Sunnyvale, CA, U.S.A.) equipped with a CAMCORDER GZ-EX210 (JVC, Bad Vilbel, Germany) was used for the optical observations. The same heating rate of 5 K/min was employed, but without inert gas protection. Screen shot images were extracted from continuous videos registered from 160 to 320 °C, for all amino acids significant moments are shown in section 3.

## 3. Experimental results

### 3.1 Data

Although we examined all 20 amino acids, we report results for those eight of them, for which the sum of the volatile gases, NH_3_, H_2_O, CO_2_ and H_2_S, matched the mass loss registered by thermogravimetry (TG). For each amino acid we show the skeleton structure, the optical observations, the DSC signal in red and the TG signal in black, as well as the ion currents for important channels, quantitatively significant are only the 17 Da (NH3, green lines), 18 Da (H2O, blue lines), and 44 Da (CO2, grey lines) signals. The logarithmic scale overemphasizes the molecular weights. The DSC data are given in W/g, the TG data in percent. The QMS data are ion currents [A] per sample.

**Figure 3.1.1.**
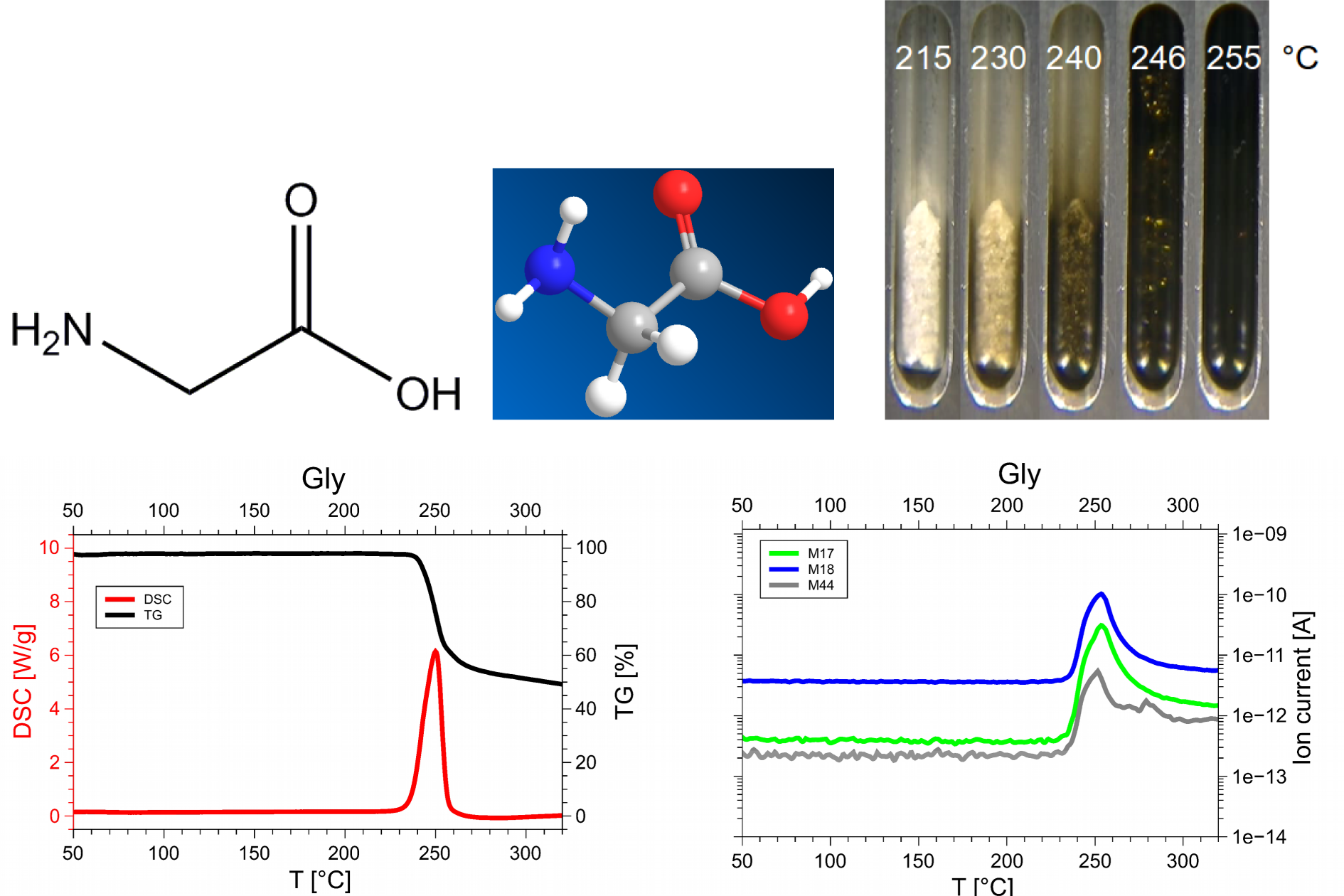
Glycine, Gly, G, C2H5NO2, 75 Da, Hf = −528 kJ/mol

**Figure 3.1.2.**
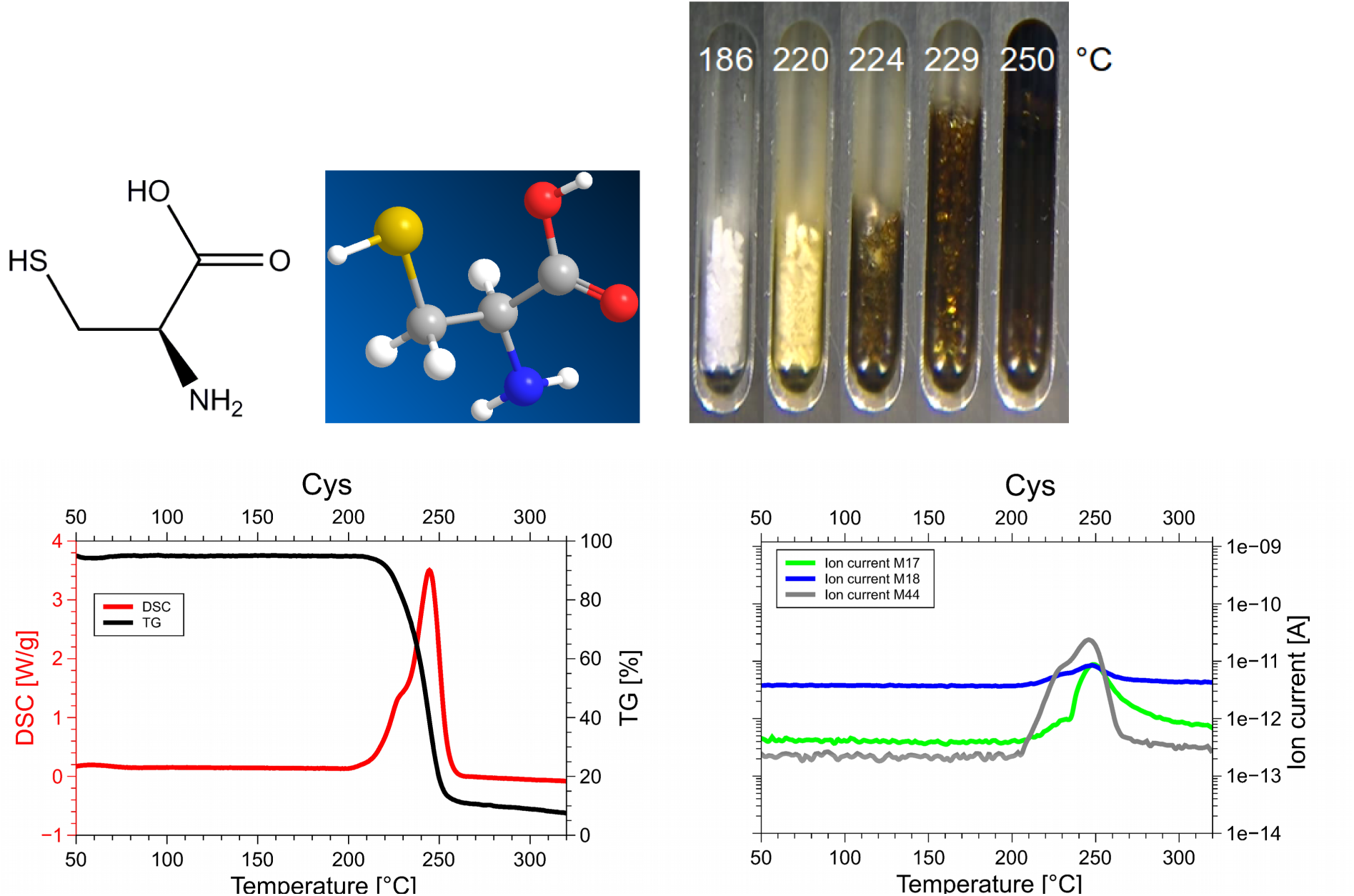
Cysteine, Cys, C, C3H7NO2S: 121 Da, Hf = −534 kJ/mol

**Figure 3.1.3.**
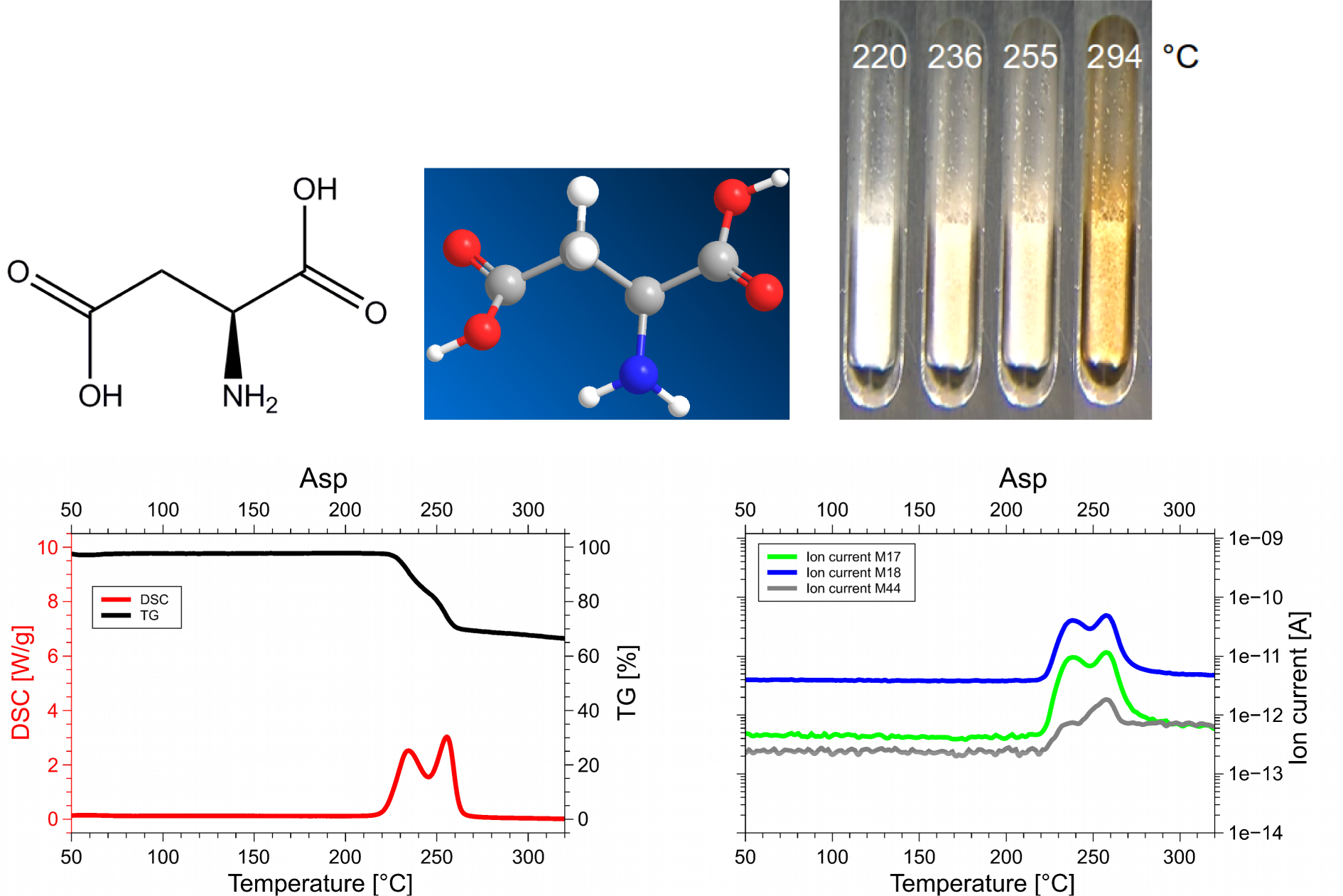
Aspartic acid, Asp, D, C4H7NO4: 133 Da, Hf = −973 kJ/ mol

**Figure 3.1.4.**
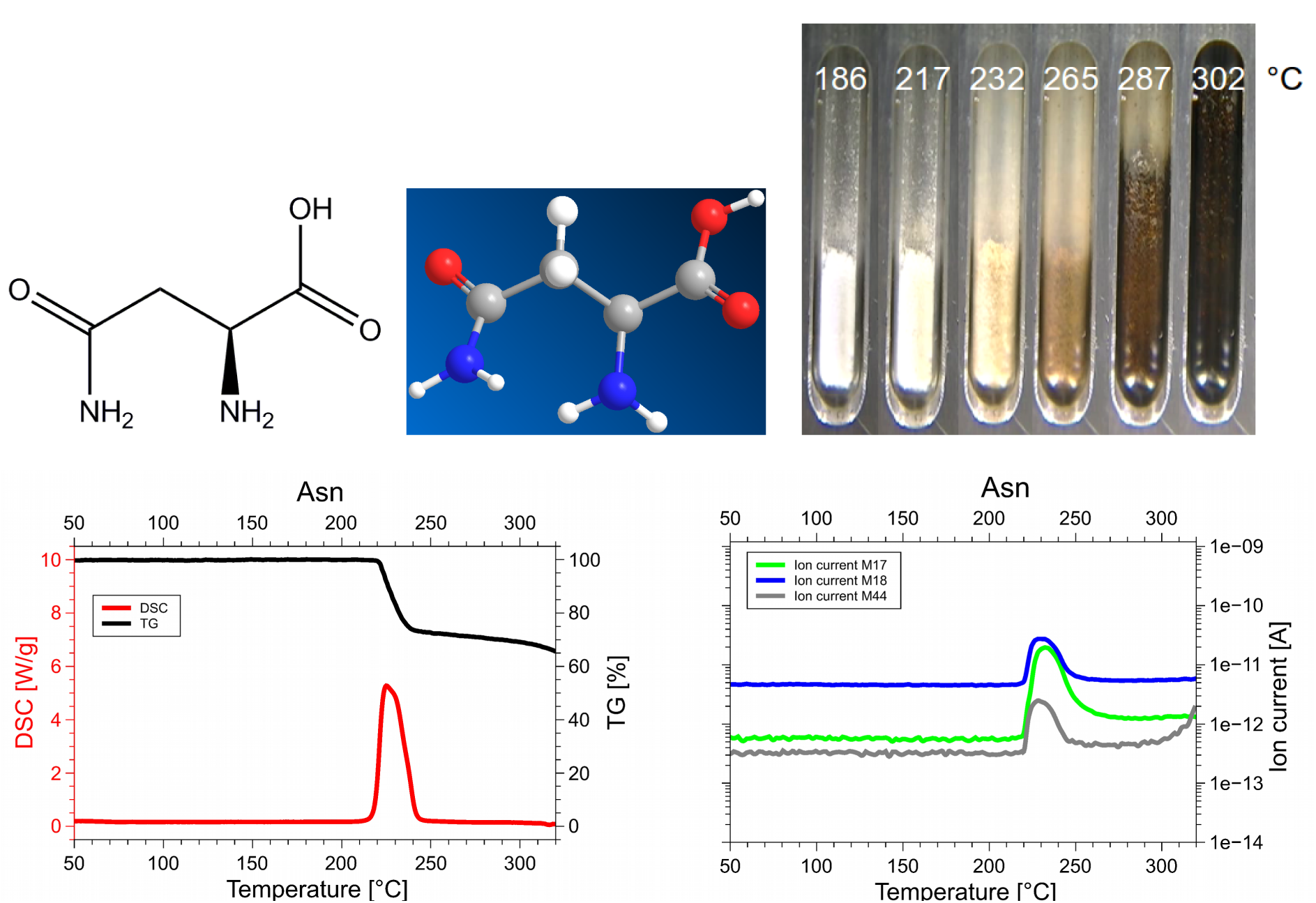
Asparagine, Asn, N, C4H8N2O3: 132 Da, Hf = −789 kJ/mol

**Figure 3.1.5.**
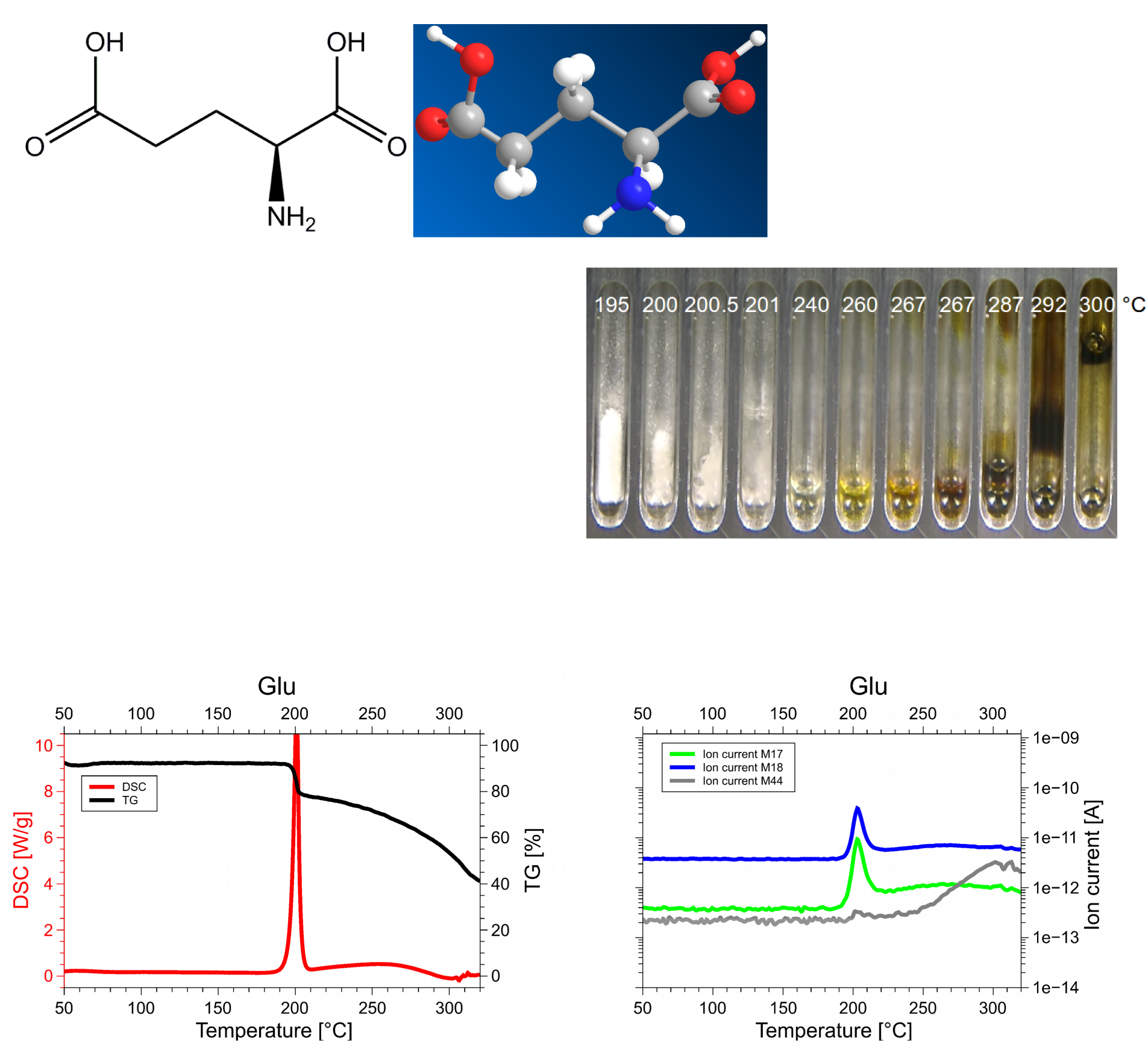
Glutamic acid, Glu, E, C5H9NO4: 147 Da, Hf = −1097 kJ/mol

**Figure 3.1.6.**
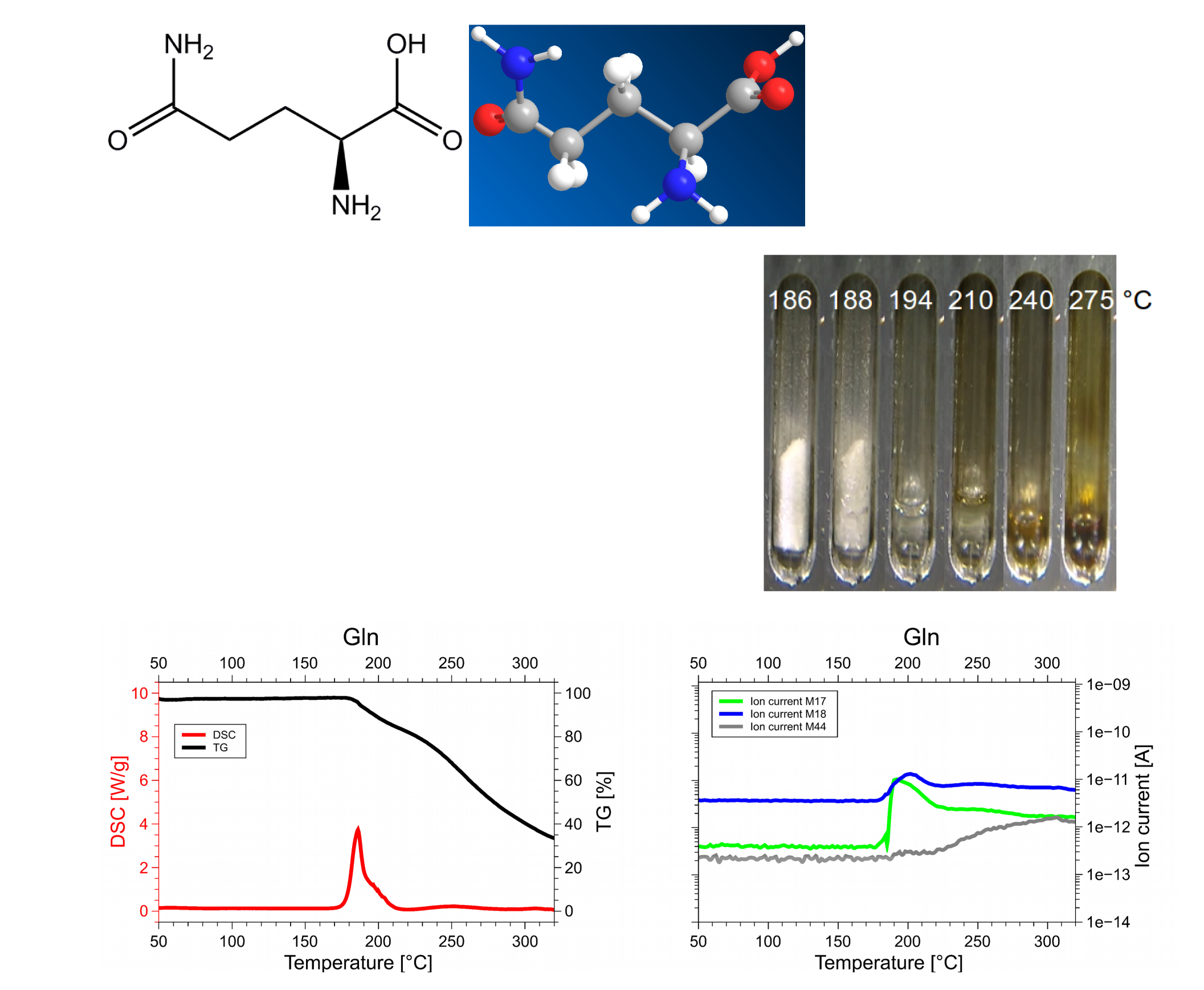
Glutamine, Gln, Q, C5H10N2O3: 146 Da, Hf = −826 kJ/mol

**Figure 3.1.7.**
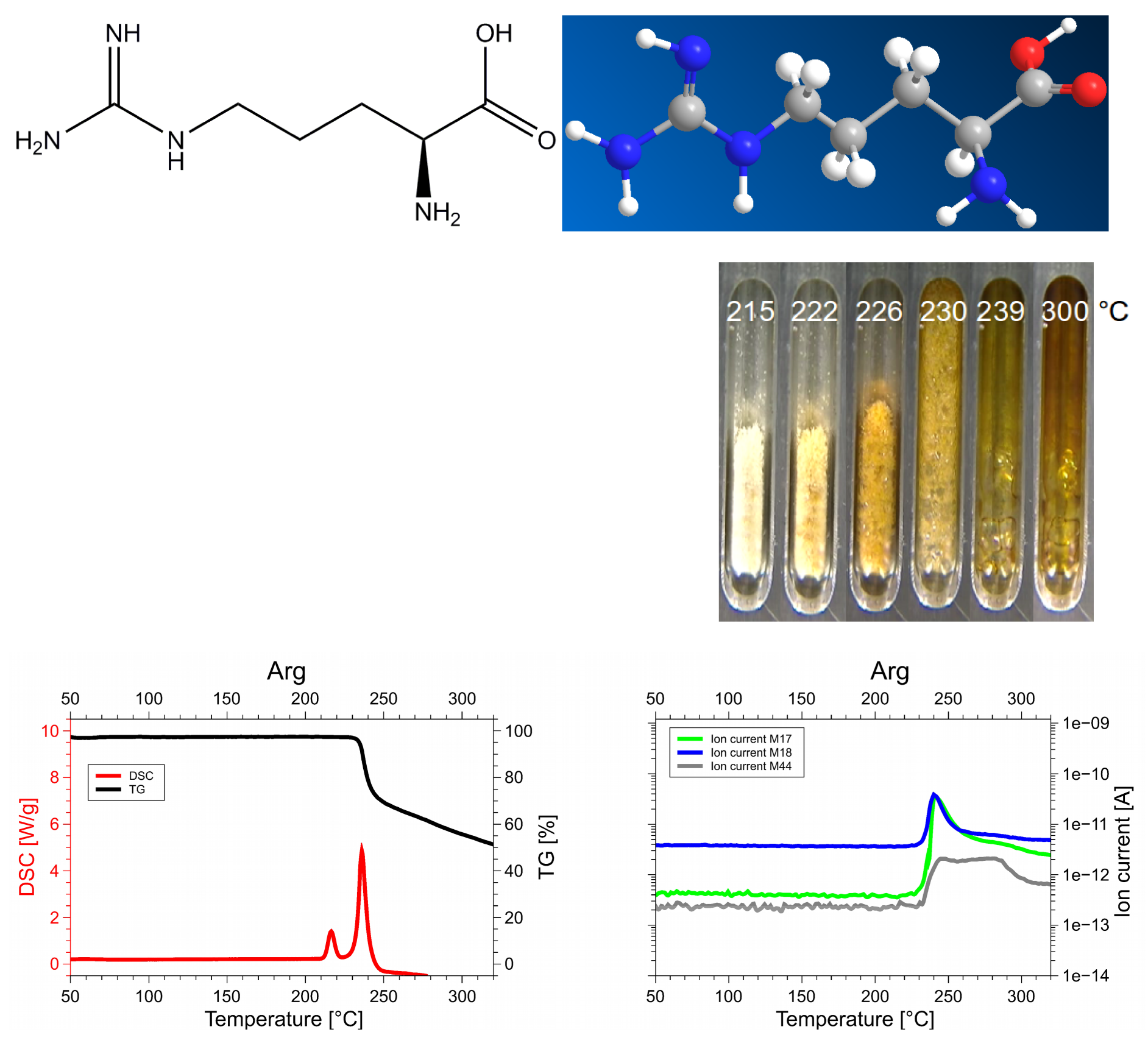
Arginine, Arg, R, C_6_H_14_N_4_O_2_: 174 Da, H_f_ = −623 kJ/mol

**Figure 3.1.8.**
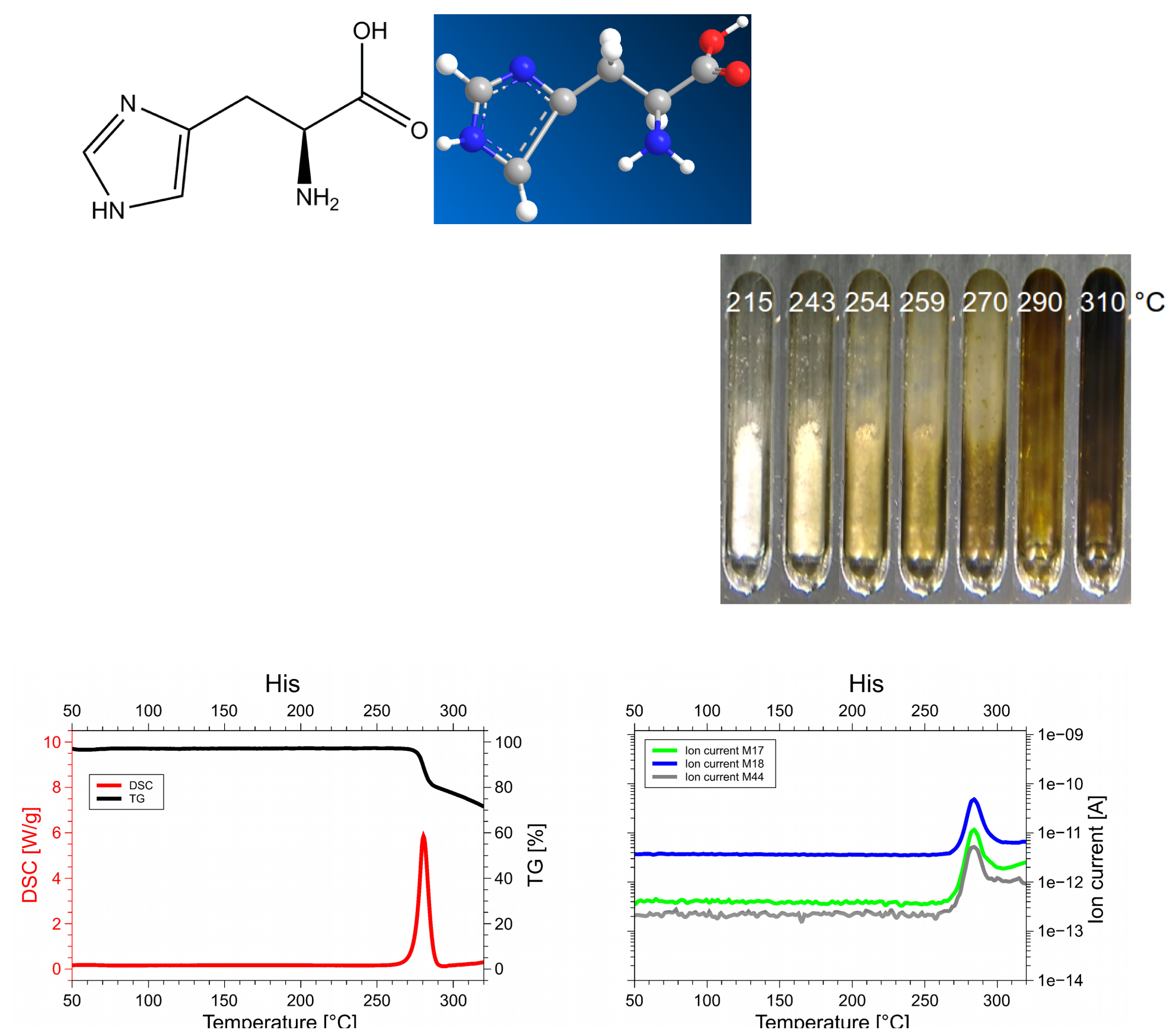
Histidine, His, H C6H9N3O2: 155 Da Hf = −467 kJ/mol

The DSC, TGA and QMS curves share one essential feature: In DSC there are peaks at a certain T_peak_ temperature for each amino acid, at the same temperatures they are accompanied by drops in TGA and QMS peaks. The simple fact that the DSC and QMS signals coincide in bell shaped peaks with the TGA drop proves that essentially one simple decomposition process takes place, there is not a spectrum of decomposition temperatures, as there would be for proteins. Qualitatively this proves that the process observed is neither melting nor sublimation (as claimed in the literature, [Acree and Chickos, 2010]). The observed process is decomposition, none of the eight amino acids exists in liquid form.

The optical observations, not obtained under vacuum but under some air access, are informative nevertheless. Solid/liquid transitions, with the liquid boiling heavily, coincide with the peak temperatures for Gly, Cys, Gln, Glu, Arg and His. Only for Asn and Asp there are solid/solid transformations at the peak temperatures. For Asn there is liquification at 280°C, Asp stays solid up to 320°C.

The DSC signals have the dimension of specific power [W/g], the QMS ones are ion currents of the order of pA. Integration over time, or, equivalently, temperature, gives the peak areas, which are specific energies [J/g] and ionic charges, of the order of pC. Reduction from specimen weights, typically 10 mg, to mol values is trivial. In absolute terms the ion currents and ionic charges are meaningless, because equipment dependent, calibration is needed. Only one reliable calibration substance was available, sodium bicarbonate (NaHCO_3_) = X1. It decomposes upon heating,

2 NaHCO_3_ ^ ⟶Na_2_CO_3_ + CO_2_ + H_2_O. The ^ CO2 mol/mol NaHCO_3_ and ^ H_2_O mol/mol NaHCO_3_ lines were quantitatively repeatable over months, in terms of pC/mol CO_2_ and pC/mol H2O. They served to identify 1 mol CO_2_/mol Cys and ^ mol H_2_O/mol Q beyond any doubt. In the absence of primary NH_3_ calibration we had to resort to secondary substances, glutamine, aspartic acid and asparagine, which retained stable NH_3_ and H_2_O signals over months. The ^ mol NH_3_/mol Q can only come from the glutamine dimer, which implies that also the H_2_O signal from glutamine corresponds to ^ mol H_2_O/mol Q. For the other two, the correspondence between 1 mol H_2_O and 1 mol NH_3_ is convincing. Thus we had four consistent reference points: mol H_2_O and mol CO_2_ from NaHCO_3_, and mol NH_3_ from glutamine, and 1 mol NH_3_ from Asparagine.

For each amino acid sample, the ion current is measured individually in each mass channel between 1 and 100 Da in 1 Da intervals. Integration over time (and temperature) gives for each mass the ion charge per mol AA, [C/molAA], and with the four calibrations the final values of mol/molAA. In Figs. 3.2 to 3.4 the ion charges are plotted on the left, the mol amounts on the right. In the graph for 17 Da (Figure 3.2) there appeared a 20 μC/mol signal for the reference substance X1. Since this definitely cannot contain NH_3_, a systematic error of 20 μC/mol must be present, though the statistical errors are smaller.

**Figure 3.2.**
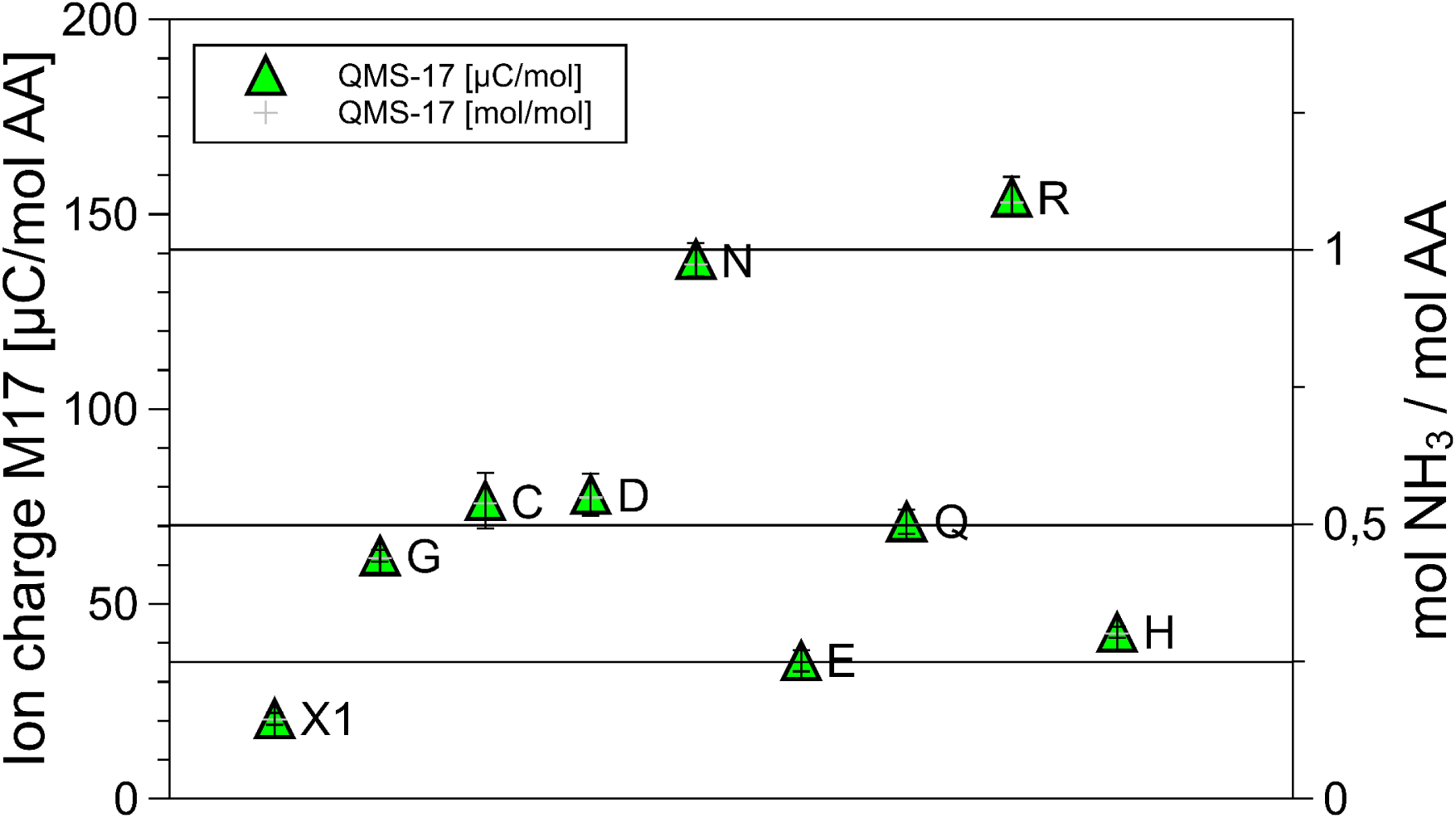
Signals in the 17 Da, the NH_3_ channel, for each of the amino acids. Ionic charges in the peaks on the left, mol NH_3_/mol amino acid on the right. The clustering of G, C, D, Q around mol NH_3_/mol AA and of N and R around 1 mol NH_3_/mol AA is striking.

**Figure 3.3.**
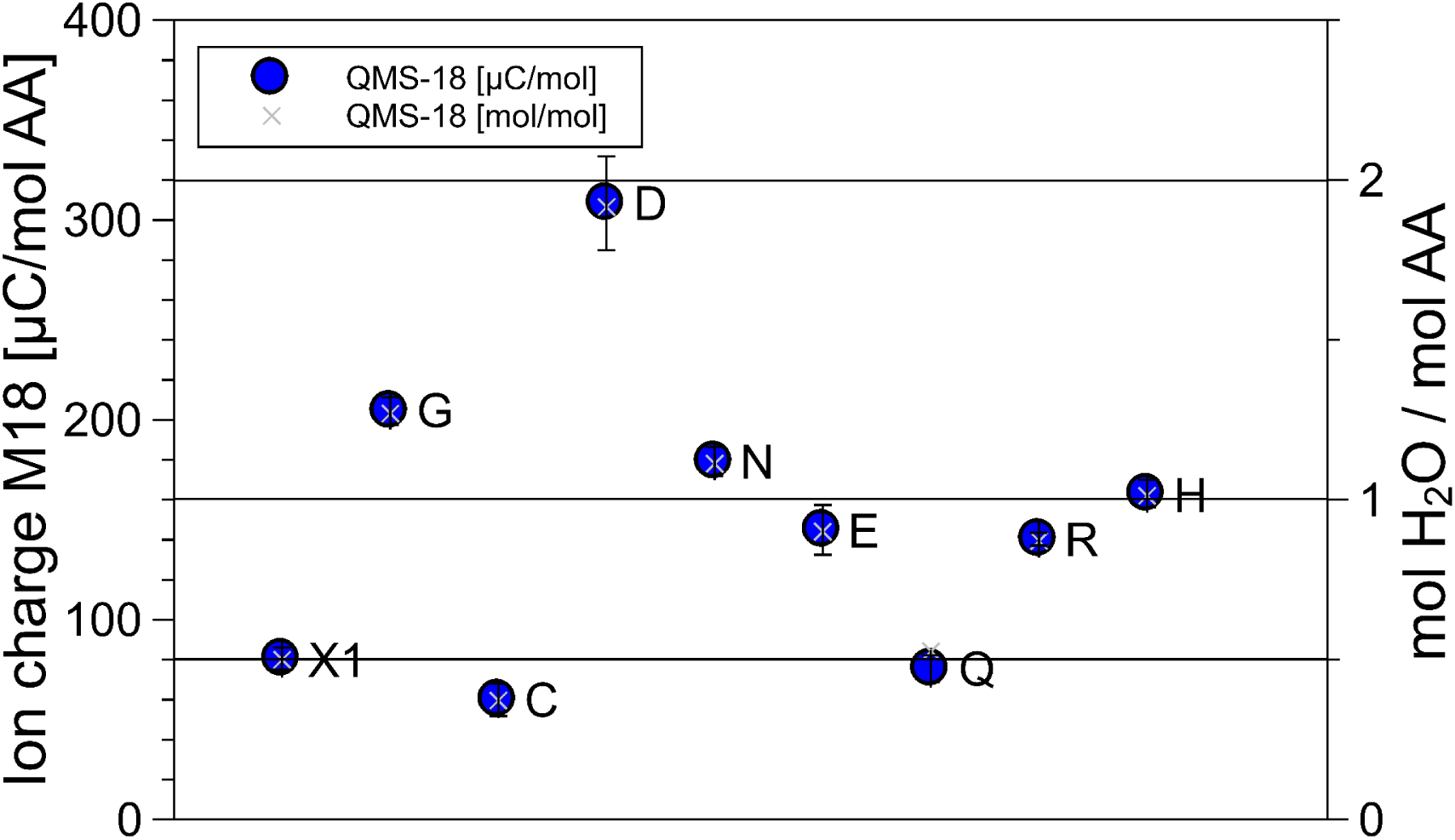
Signals in the 18 Da, the H_2_O channel, for each of the amino acids. Ionic charges in the peaks on the left, mol H_2_O/mol amino acid on the right. The clustering of C and Q around the mol H_2_O level, of N, E, R, H around the 1 mol H_2_O level, and the 2 mol point for D are striking.

**Figure 3.4.**
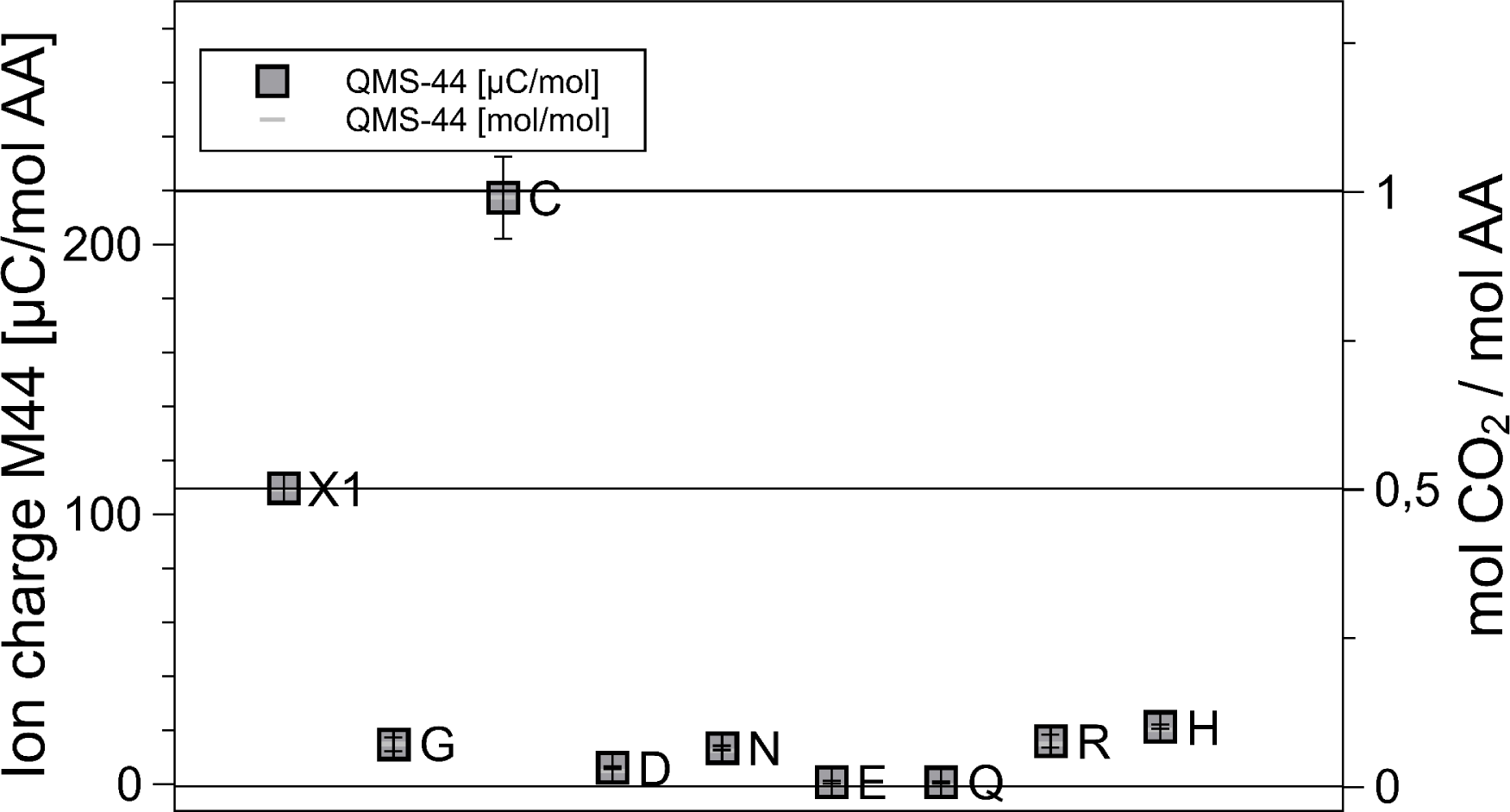
Signals in the 44 Da, the CO_2_ channel, for each of the amino acids. Ionic charges in the peaks on the left, mol CO_2_/mol amino acid on the right. Only C produces 1 mol CO_2_, the level of the others is negligible.

The absolute ionic currents of Figures 3.2 to 3.4 are equipment dependent and not significant, but the relative values are encouraging. One mol NH_3_ produces 12% less and CO_2_ 54 ***%*** more ions than one mol H_2_O. Indeed the ionization cross sections of NH_3_, H_2_O and CO_2_ are reported to be in that order [NIST website, Kim et al.].

**Table 1.** The first two columns show the names of the amino acids. The third is the heat of formation, H_f_(s) of the solid amino acid. The other columns are experimental data. T_peak_ is the temperature of the DSC and QMS peaks, H_peak_ is the endothermic peak area, ΔM the mass loss registered by TGA. QMS-17, QMS-18 and QMS-44 are the mol fraction found in QMS for the 17 Da, 18 Da and 44 Da peaks, respectively. From these mol fractions the sum of the gaseous mass, M(gas), is calculated. The last column, AΔM-M(gas), is the difference between the mass loss ΔM measured in TGA, and the mass loss M(gas) calculated from QMS. If there are a, b and c mols of NH_3_, H_2_O and CO2, respectively, M(gas) = 17a+18b+44c. The small values of ΔM-M(gas) prove that no gases other than NH_3_, H_2_O and CO_2_ evolved (For Cys the mol H_2_S, 34 Da, has been added). The enthalpy of formation of the residue is calculated as H_res_ = H_f_ – 45a – 242b – 396c.

| AAA | A | M.W.<br>Da | H<sub>f</sub><br>kJ/mol | T peak<br>°C | H peak<br>kJ/mol | Δ M<br>Da | QMS-17 |  | QMS-18 |  | QMS-44 |  | M (gas)<br>Da | Δ M - M (gas)<br>Da | H res<br>kJ/mol |
| --- | --- | --- | --- | --- | --- | --- | --- | --- | --- | --- | --- | --- | --- | --- | --- |
| Gly | G | 75 | -529 | 250 | 64 | 35 | 0,44 ± | 0,01 | 1,27 ± | 0,04 | 0,07 ± | 0,00 | 34 | 1 | -172 |
| Cys | C | 121 | -534 | 245 | 94 | 98 | 0,54 ± | 0,05 | 0,37 ± | 0,05 | 0,99 ± | 0,07 | 59 | 39 | -27 |
| Asn | N | 132 | -789 | 225 | 118 | 35 | 0,97 ± | 0,03 | 1,11 ± | 0,05 | 0,06 ± | 0,00 | 39 | -4 | -451 |
| Gln | Q | 146 | -826 | 185 | 77 | 22 | 0,50 ± | 0,02 | 0,53 ± | 0,12 | 0,00 ± | 0,00 | 18 | 4 | -675 |
| Asp | D | 133 | -973 | 256 | 121 | 39 | 0,55 ± | 0,04 | 1,92 ± | 0,15 | 0,03 ± | 0,00 | 45 | -6 | -473 |
| Glu | E | 147 | -1010 | 201 | 89 | 20 | 0,25 ± | 0,02 | 0,90 ± | 0,08 | 0,00 ± | 0,00 | 21 | -1 | -779 |
| Arg | R | 174 | -624 | 236 | 54 | 47 | 1,09 ± | 0,04 | 0,87 ± | 0,02 | 0,07 ± | 0,01 | 37 | 9 | -334 |
| His | H | 155 | -467 | 280 | 82 | 27 | 0,30 ± | 0,01 | 1,01 ± | 0,04 | 0,10 ± | 0,00 | 28 | -1 | -169 |

**Figure 3.5.**
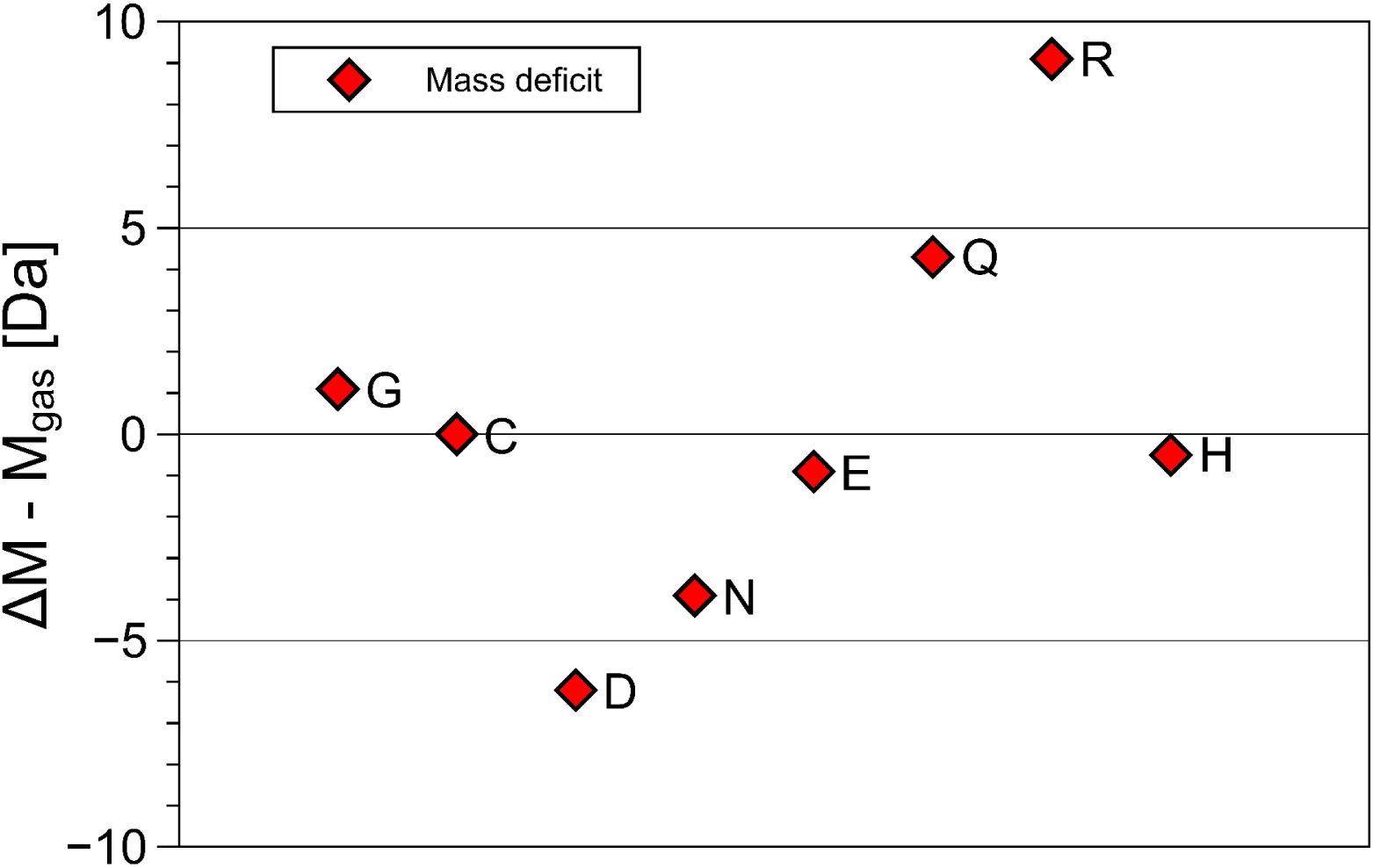
The difference between the mass loss registered by TGA, ΔM and the volatile mass found as NH_3_, H_2_O, CO_2_ and H_2_S, ΔM - Mgas remains below |9| Da. This is confirmation that no other gases are produced.

Figures 3.2 to 3.5 and Table I summarize the experimental data: with the exception of cysteine, thermal decomposition results in three gases, mainly H_2_O, less NH_3_ and hardly any CO_2_. The weight of these three gases adds up to the weight loss registered by TGA, therefore no other gases evolve in appreciable amount - they are not seen in QMS either. The proximity of the molfractions to integer or half-integer values indicates simple decomposition chains. The process causing the peaks cannot be melting (because of the mass loss), nor sublimation (because of the QMS signals). One concludes that amino acids do not exist in liquid or gaseous form. They decompose endothermally, with heats of decomposition between −72 and −151 kJ/mol, at well defined temperatures between 185°C and 280°C.

### 3.2 General remarks

These amino acids consist of different side chains attached to the Ca of the same backbone, NH_2_-Ca-(C*OOH), but their decomposition chains are quite different. The pyrolytic process is controlled by three balance laws: In terms of Da the masses must add up, chemically the atomic species must balance, and the enthalpy of formation must equal the enthalpies of formation of the products plus the endothermic heat of reaction. The amounts of volatile products are experimental values (TGA and QMS). For the residues only the mass is experimental, their composition is inferred. In section 4 we analyze possible pathways. Although the choices, restricted by compositional, mass and enthalpy considerations, are convincing, they cannot be unique beyond doubt. Alternatives to our proposals, but indistinguishable by us, are possible. Analyses of the decomposition chains are, therefore, tentative or speculative. Nevertheless, they are less speculative than those of Rodante et al. (1992), who had only TGA and DSC, but no QMS at their disposal. What Acree and Chickos (2010)call “sublimation enthalpies” agree more or less with our decomposition enthalpies. One concludes that they must refer to decomposition, not merely sublimation without composition change.

We use of the enthalpy values listed for standard conditions (Haynes 2013) without minor corrections for specific heats and entropies up to the actual reaction temperatures. Moreover, hydrogen gas escapes our attention, it is too light (2 Da) to be registered in QMS and TGA, nor does it appear in the enthalpy sum, its heat of formation being zero by definition. With exception of hydrogen, the mass balance, controlled by TGA, confirms that beyond the residue and the three gases, nothing else is formed. The real constraint is the enthalpy balance. For the enthalpy balance production of water is necessary. The expression of the formation enthalpies of the 20 amino acids CaHbNcOdSe has the least square fit Hf(CaHbNcOdSe) = 30.3a - 37.8n +16.5c - 182.4d - 71.3e [kJ/mol]. The oxygens outweigh the others with −182 kJ/mol. The obvious way of efficiently transferring enthalpy from the reactants to the products is the formation of water, with H_f_(H_2_O) = −242 kJ/mol.

Detailed analysis for each amino acid is helped by preliminary reference to a few reactions possible in principle. CO_2_ production in Cys is obviously a special case. In principle one expects the N-termini to be stable, making desamination to produce NH_3_ unlikely. Nitrogen in the side chains is another matter, indeed the NH_3_ producing Asn and Arg have nitrogen in their side chains. The predominance of H_2_O production indicates instability of the C-terminus beyond the C* atom, where dehydration can occur by n-oligomerization, which yields (n-1)/n mol H_2_O/mol AS, from dimerization for n=2, to 1 mol H_2_O/mol for n ^ ~ in polymerization. A special case of dimerization is external cyclization in the diketopiperazine reaction, which yields 1 mol H_2_O/mol AA. These involve joining N– and C-termini in a dehydration reaction. For long side chains also internal cyclization, where the end of the side chain connects to the C-terminus can be envisaged. Integer and half-integer mol values restrict the choice for the residues, but not unequivocally.

## 4 Data analysis, amino acid by amino acid

All DSC peaks are endothermic, their areas are given negative signs. With this convention endothermic evaporation and exothermic production of water are written as

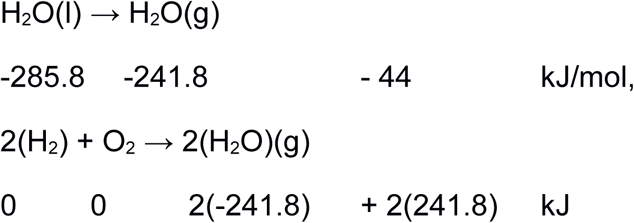

### 4.1 Glycine

Glycine, Gly, G, C_2_H_5_NO_2_, 75 Da, H_f_ = −528 kJ/mol.

Simple endothermic peak at 250°C, H_peak_ = −72.1 kJ/mol.

The QMS signal of 3/2 mol H_2_O/mol Gly plus % mol NH_3_/mol Gly is beyond doubt, it is confirmed by the mass loss of 35 Da/mol Gly. This leaves only 10% of the original hydrogen for the residue. The triple and double bonds in carbon rich C_4_HNO, C_3_HNO and C_2_HNO preclude them enthalpy wise and make deposition of carbon likely,

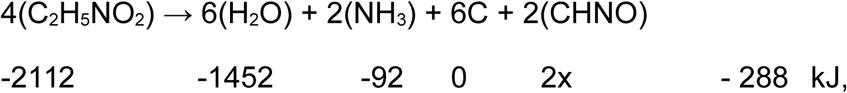

leaving −280 kJ/mol = x for the moiety CHNO, which is the composition of the peptide bond. Chemspider lists two symmetric molecules consisting entirely of peptide bonds, 1,3-Diazetine-2,4-dione, C_2_H_2_N_2_O_2_, chemspider 11593418, 86 Da, (Figure 4.1a) or its isomer 1,2-Diazetine-3,4-dione, C_2_H_2_N_2_O_2_, chemspider 11383421, 86 Da, Fig 4.1b. The scarcity of hydrogen is such that not even the smallest lactam, 2-Aziridinone, C_2_H_3_NO, 57 Da, chemspider 10574050, bp 57 °C (Figure 4.1c) can serve as residue.

**Figure 4.1a.**
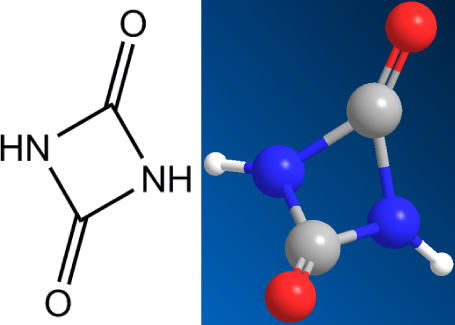
Residue of Gly, C_2_H_2_N_2_O_2_, 1,3-Diazetine-2,4,dione, 86 Da

**Figure 4.1b.**
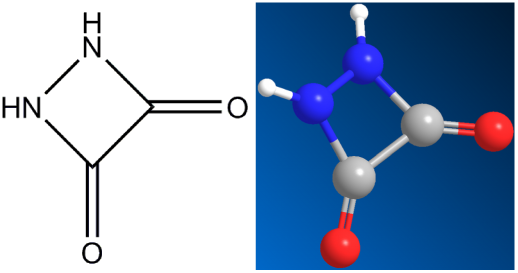
Isomer of 4.1a, 1,2-Diazetine-3,4-dione, 86 Da

**Figure 4.1c.**
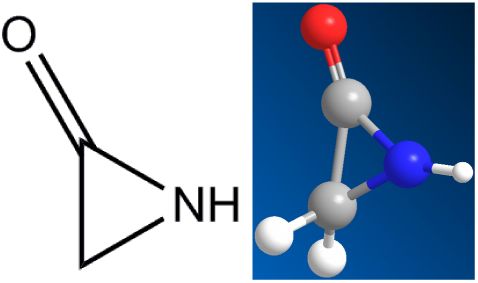
2-Aziridinone, C_2_H_3_NO, 57 Da

**Figure 4.1d.**
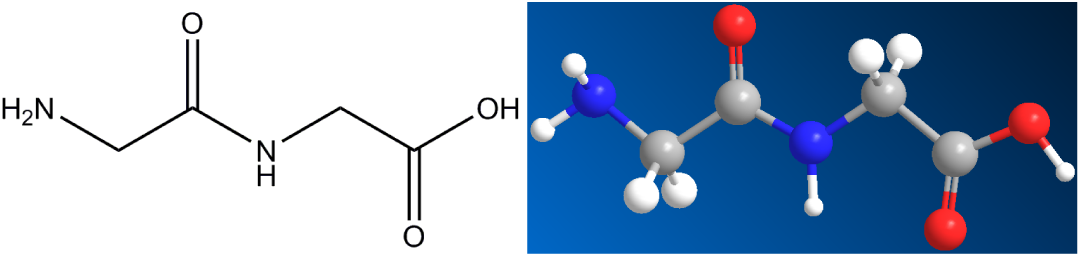
Intermediate dimer, glygly, C_4_H_8_N_2_O_3_, 132 Da.

The simplest pathway seems the formation of linear Glycylglycine, chemspider 10690 (Figure 4.1d), m.p. 255°C (which is T_peak_!) C_4_H_8_N_2_O_3_, H_f_(s) = −748 kJ/mol [Haynes 2013] from which the central peptide bond-(C=O)–NH– is detached by cutting off the NH_2_–Cα–H2– group on one, and the Cα–H_2_–C*OOH group on the other side. The former makes NH3 plus C, the latter makes 2C plus 2(H2O). This process is specific for the glycine dimer, in which the Cα atoms are not protected by proper sidechains, they are just –Cα–H2– units. This pathway to shear peptide bonds is of interest in the context of possible peptide nucleic acid (PNA) synthesis [Banack et al. 2012] via N-2– aminoethylglycine (AEG), C_4_H_10_N_2_O_2_, chemspider 379422, which is deoxidized diglycine, 2Gly ^ O2 + AEG.

### 4.2 Cysteine

Cysteine, Cys, C, C_3_H_7_NO_2_S: 121 Da, H_f_ = −534 kJ/mol.

T_peak_ = 221°C with a mass loss of 98 Da, H_peak_ = −96 kJ/mol.

The clear 1 mol CO2 signal leaves no oxygen to form H_2_O, therefore the spurious 18 Da line must stem from a systematic error. There is also % mol NH3/molCys. For H_2_S there is indeed a signal at 34 Da. It corresponds to 1 mol, because the ionization cross sections of H_2_S and H_2_O are nearly identical, so that the calibration of Figure 3.3 applies. The mass loss of 44+34+8.5 = 70% of 121 Da agrees with TGA.

Chemical analysis found no sulfur in the residue. No possibility for forming disulfide bridges between molecules is left. Neither COS nor CS2 was found. The total reaction is

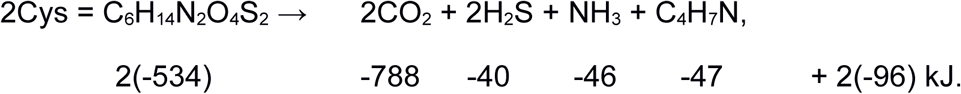

On the left −1068, on the right −1113 kJ. Pathway to the formation of C_4_H_7_N might be ejection of the carboxyl group–C*OOH and the –SH group from Cys, the remaining chain NH_2_-Caα-C* is too short for internal, but suitable for external cyclization. Two of these form the asymmetric 5-ring (3-pyrrolidinamine, 144134), Figure 4.2a, from which the –NH2 is cutt off. Indeed the % NH_3_ ejected confirms such dimerization. That leaves the molecule C_4_H_7_N: 2,5-Dihydro-1H-pyrrole, chemspider 13870958, b.p. 90°C, 69 Da, H_f_(s)=-46.6kJ/mol (Figure 4.2b), or another pyrroline, with the double bond elsewhere in the ring. Indeed there is heavy boiling beyond the peak. In view of the richness in hydrogen, several small hydrocarbon lines are not surprising.

**Figure 4.2a.**
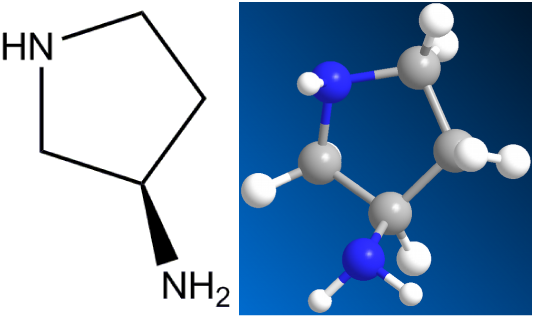
Intermediate compound: 3-pyrrolidinamine, chemspider 144134, 86 Da.

**Figure 4.2b.**
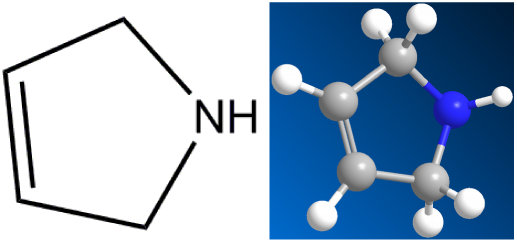
Residue of Cys, C4H7N, 2,5-Dihydro-1H-pyrrole, chemspider 13870958, 69 Da.

### 4.3 Aspartic acid

Aspartic acid, Asp, D, C_4_H_7_NO_4_: 133 Da, H_f_ = −973 kJ/ mol.

DSC shows two distinct peaks, at 230°C and at 250°C, in each of which 1 mol H_2_O/mol Asp is ejected. The endothermic heats are −64 and −61 kJ/mol, respectively. Stays powder up to 294°C, i.e. solid/solid transformation in peak. The reaction

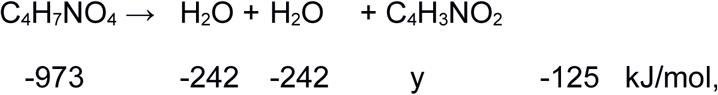

with calculated y = −364 kJ/mol, which is reasonable for the formation enthalpy of the polysuccinimide unit (PSI). The molecular weight of C_4_H_3_NO_2_ is 97 Da.

The compound (C_4_H_3_NO_2_)_n_ is polysuccinimide. The two peaks prove that the reaction occurs in two steps, in the first at 230°C the condensation reaction produces polyaspartic acid, Asp ^ H_2_O + poly-Asp, in the second at 250°C the poly-Asp degrades to polysuccinimide (PSI) by ejection of another 1 mol H_2_O/mol Asp. Such a reaction was reported by Schiff [1897]. The molecule drawn is beta-poly-Asp, there is an isomer, alpha-poly-Asp, where the next C in the ring forms a bond to its neighbor. We have no possibility to decide between the two.

**Figure 4.3.**
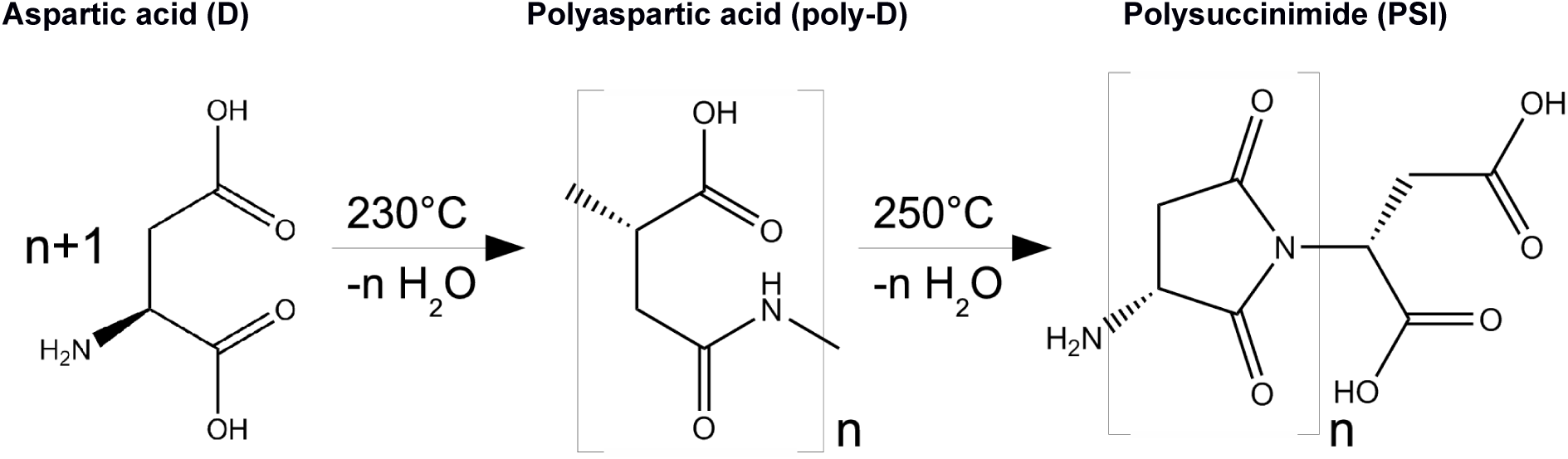
The pathway from Aspartic acid (D) to polysuccimide (PSI). Compared with succinimide, the N-C bond in polysuccinimide economizes two hydrogen atoms.

### 4.4 Asparagine

Asparagine, Asn, N, C_4_H_8_N_2_O_3_: 132 Da, H_f_ = −789 kJ/mol.

In the broad peak at 232°C, 1 mol H_2_O /mol Asn and 1 mol NH_3_ /mol Asn are ejected. H_peak_= −122 kJ/mol. The product stays a white powder up to 265°C, i.e. there is a solid/solid transformation in the reaction

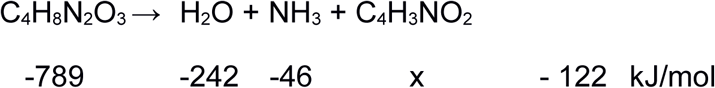

with calculated x = −379 kJ/mol. In the Asp decomposition, H_f_(C_4_H_3_NO_2_) was calculated as y = −364 kJ/mol. The two values agree, although because of their histories, the two PSI are not identical. If Asn followed the example of Asp, it would eject 1mol H_2_O /mol Asn in the condensation reaction, Asn ⇾ H_2_O + poly-N, followed by degradation of poly-N to polysuccinimide (PSI) by ejection of 1mol NH_3_ /mol Asn. If, however, the H_2_O of the condensation reaction is not ejected but retained, it can replace the –NH_2_ in poly-N by –OH. According to Asn ⇾ NH_3_ + poly-D this amounts to the formation of polyaspartic acid from asparagine by ejection of NH_3_. The poly-D then degrades to polysuccinimide(PSI) by ejection of 1mol H_2_O /mol Asn. Apparently both alternatives occur and there is one broad peak containing both NH_3_ and H_2_O.

**Figure 4.4.**
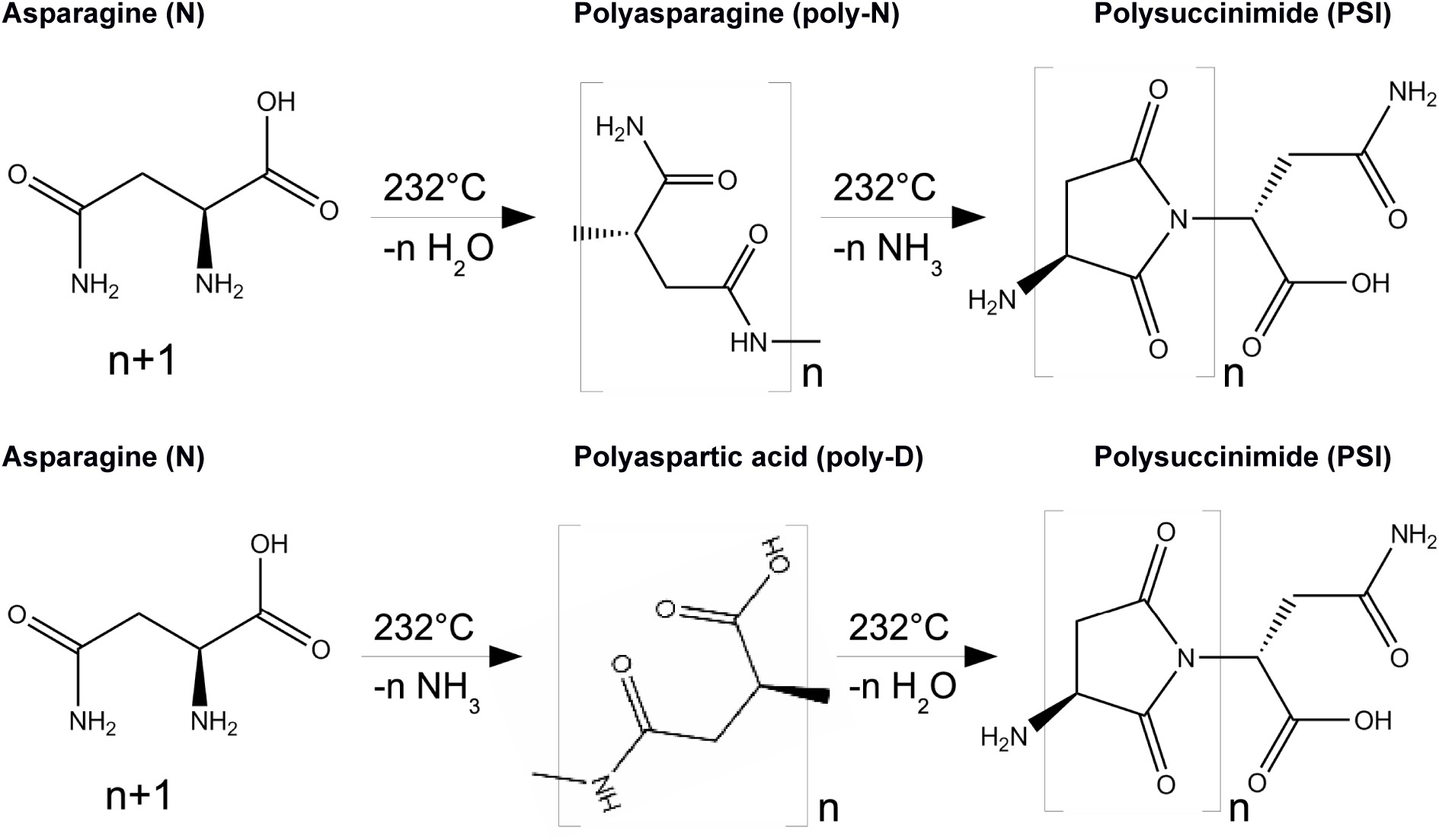
Two pathways from asparagine (N) to polysuccinimide (PSI): either through polyasparagine (poly-N) or polyaspartic acid (poly-D). Compared with succiminide, the N-C bond in polysuccinimide economizes two hydrogen atoms.

Though the formulae for PSI formed from Asp and from Asn, are the same, (C_4_H_3_NO_2_)_n,_ these two residues need not be identical. For kinetic reasons the oligomerization or polymerization might have proceeded to different lengths in polyAsp and poly-Asn, therefore the degraded products PSI might have different lengths, with different stabilities and melting points. Moreover, the telomers are different, –OH for PSI from Asp and –NH_2_ for PSI from Asn. Indeed PSI from Asp remains a white powder up to 289°C, while PSI from Asn starts melting at 289°C.

### 4.5 Glutamic acid

Glutamic acid, Glu, E, C_5_H_9_NO_4_: 147 Da, H_f_ = −1097 kJ/mol; T_peak_ = 200°C, H_peak_ = −88 kJ/mol.

At 200°C, 1 mol H_2_O /mol Glu is seen in QMS, the DSC area is −121 kJ/mol, mass loss in the peak is 12% (17 Da). The dehydration of Glu has been known for a long time [Haitinger, L., Vorlaeufige Mittheilung uber Glutaminsaure und Pyrrol. Monatsh. Chem., 1882, 3, 228–229].

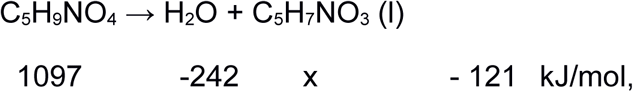

with calculated x = −734 kJ/mol for H_f_(C_5_H_7_NO_3_), pyroglutamic acid, chemspider 485, 129 Da, T_m_ = 184°C, b.p. = 433°C.

**Figure 4.5a.**
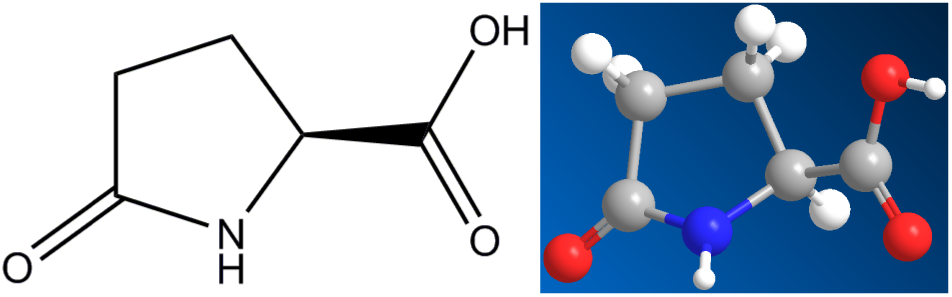
The final residue of Glu, pyroglutamic acid, C_5_H_7_NO_3_, 129 Da

This lactam is biologically important, but its enthalpy of formation is apparently not known. Known is the H_f_(s) = −459 kJ/mol and H_f_(g) = −375 kJ/mol for C_4_H_5_NO_2_, succinimide, 99 Da, the five ring with O= and =O as wings (the structure is like pyroglutamic acid, but with the carboxyl group-COOH replaced by =O).

**Figure 4.5b.**
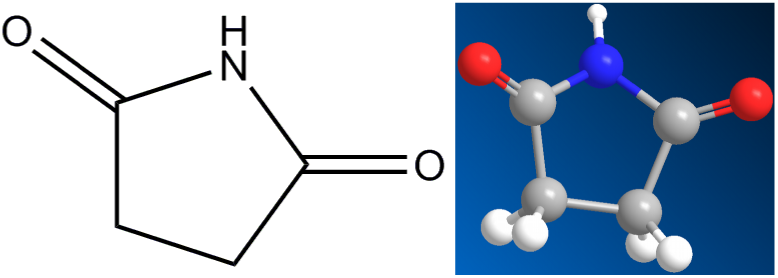
Succinimide, C_4_H_5_NO_2_, 99 Da.

The additional O should add about −200 kJ/mol, which makes the −734 kJ/mol for pyroglutamic acid plausible. The TGA weight loss beyond the peak is evaporation. Pyroglutamic acid is formed by inner cyclization of E: after the –OH hanging on Cδ_5_ is ejected, the C§δ joins the –NH_2_ hanging on Caα. This was suggested by Mosqueira et al. [2008], ours is the first experimental evidence for this process.

Since QMS does not show any CO2, the reaction to C_4_H_7_NO, the lactam pyrrolidone, 85 Da, T_m_ = 25°C, b.p. = 245°C, H_f_(l) = −286 kJ/mol, yellow liquid, can be ruled out, although cutting off the CO_2_ is sterically tempting.

**Figure 4.5c.**
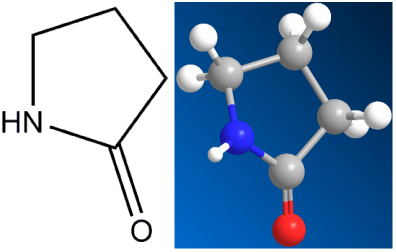
Pyrrolidone, C_4_H_7_NO, 85 Da.

### 4.6 Glutamine

Glutamine, Gln, Q, C_5_H_10_N_2_O_3_: 146 Da, H_f_ = −826 kJ/mol.

The precise half mol fractions of H_2_O and NH_3_ in the peak at T_peak_ = 185°C, H_peak_ = −77 kJ/mol, indicate that a dimer serves as intermediate step, gamma-glutamylglutamine, C_10_H_17_N_3_O_6_, chemspider 133013, b.p. 596°C:

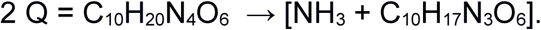

**Figure 4.6a.**
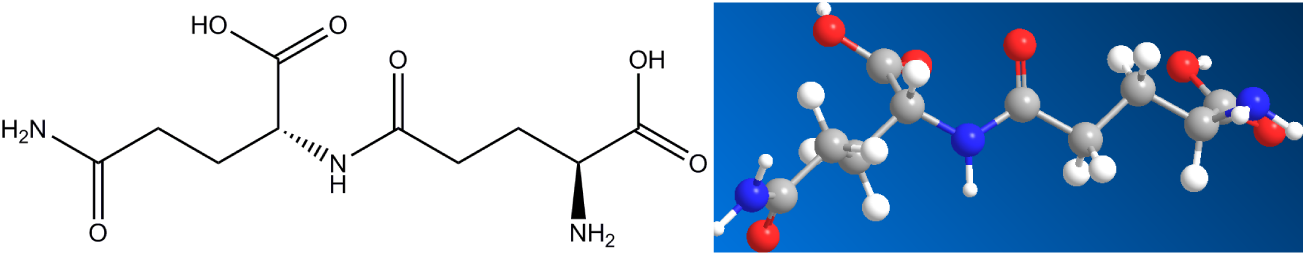
Intermediate step, gamma-glutamylglutamine, C_10_H_17_N_3_O_6_, 275 Da., After further ejection of H_2_O the total reaction is 2 Q = 2 (C_5_H_10_N_2_O_3_) —→ NH_3_ + H_2_O + C_10_H_15_N_3_O_5_.

2 Q = C_10_H_20_N_4_O_6_ → [NH_3_ + C_10_H_17_N_3_O_6_].

Chemspider lists for the residue a suitable molecule, 9185807, 5-Oxo-L-prolyl-L-glutamine, C10H15N3O5, 257 Da, b.p. 817°C, H_vap_ = 129 kJ/mol.

**Figure 4.6b.**
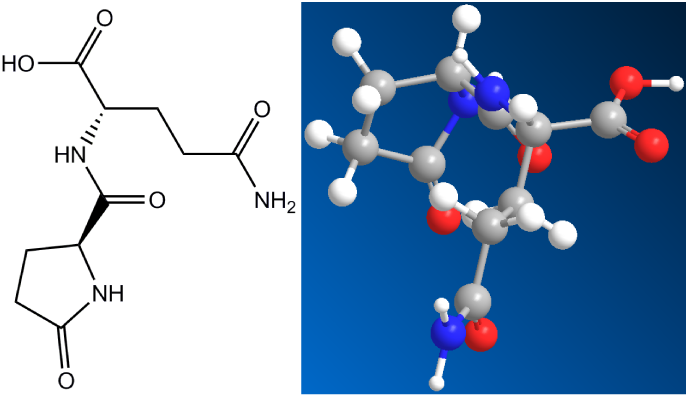
The residue of Gln: 5-Oxo-L-prolyl-L-glutamine, C_10_H_15_N_3_O_5_, 257 Da.

Above the peak at 185°C optical observations show indeed a nonboiling liquid, agreeing with the high boiling point quoted.

### 4.7 Arginine

Arginine, Arg, R: C_6_H_14_N_4_O_2_: 174 Da, H_f_ = −623 kJ/mol.

A small peak without mass loss at 220°C, −14 kJ/mol, and a main peak at 230°C, −52 kJ/mol, producing 1 mol NH_3_ plus 1 mol H_2_O in QMS, confirmed by the weight loss of 20% of 174 Da in TGA. The precursor peak without mass loss at 220°C, −14 kJ/mol, probably comes from a rearrangement in the guanidine star. In the large peak a double internal cyclization occurs. The loss of the amino group –NH_2_ in the backbone, and internal cycling joining the N next to the Cδ5 in the side chain to Caα, C_6_H_14_N_4_O_2_ ^ NH_3_ + C_6_H_11_N_3_O2, forms an intermediate, 1-Carbamimidoylproline, 157 Da, chemspider 478133.

**Figure 4.7a.**
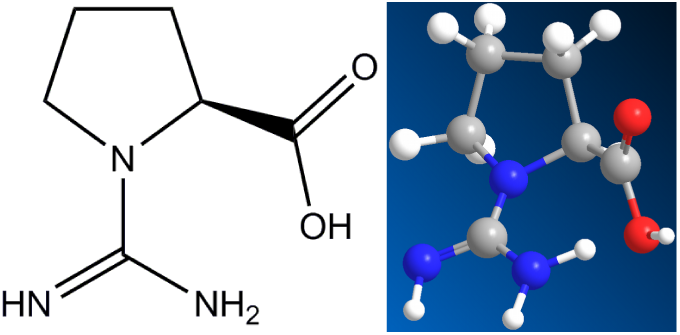
1-Carbamimidoylproline, 157 Da, representing the intermediate step after ejection of NH_3_ from Arg.

It is called “.proline”, because the ring is spanned between an N and Caα, though the N is not from the backbone. By losing the –OH and a second inner cyclization joining the =NH or the –NH2 to C*, one or the other tautomer of the final residue is formed. The total reaction is C_6_H_14_N_4_O_2_ ^ NH_3_ + H_2_O + C_6_H_9_N_3_O, not quoted in chemspider:

**Figure 4.7b.**
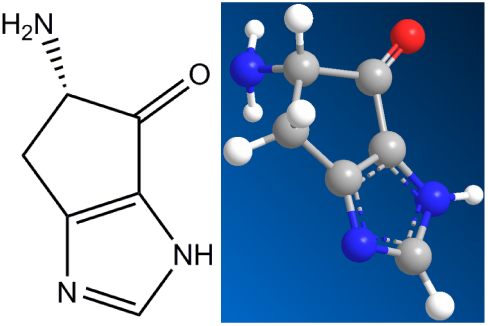
The final residue of Arg, C_6_H_9_N_3_O, 139 Da, “creatine-proline”. The creatine ring on top joins the proline ring.

This final residue is remarkable. It contains the proline ring, the guanidine star and a peptide bond in the ring of creatinine, which is the 5-ring with the =O and =OH double bonds. Creatinine, H_f_ = −240 kJ/mol, m.p. 300°C, C_4_H_7_N_3_O, chemspider 568, has several tautomeric forms. The end product in question might contain either of those rings. We have no way to decide between the alternatives, but a double ring structure seems likely.

### 4.8 Histidine

Histidine, His, H, C_6_H_9_N_3_O_2_: 155 Da, H_f_ = −466 kJ/mol.

The QMS results are clear, His ejects 1 mol H_2_O in the reaction

His = C_6_H_9_N_3_O_2_ ^ 1 H_2_O + C_6_H_7_N_3_O.

The observed 1mol H_2_O /mol His, confirmed by the weight loss of 13% of 155 Da, could stem from the condensation reaction of polymerization, but the volatility seen optically contradicts this option.

Inner cyclization seems likely. If the C* of the backbone joins the C of the imidazole ring, with =O and –NH_2_ attached outside, the 5-ring formed joins the 5-ring of the imidazole. The proposed structure would be

**Figure 4.8.**
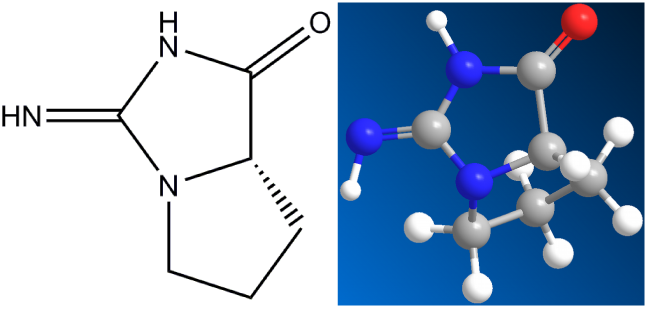
Final residue of His, C_6_H_9_N_3_O, 139 Da, consisting of two 5-rings: 2-amino-2,4-cyclopentadien-1-one (C_5_H_5_NO, chemspider 28719770) and imidazole.

Molport quotes this structure, but with the pyrazole ring (where the two N are nearest neighbours) instead of the imidazole ring (where the two N are next nearest neighbours): 5-amino-4H,5H,6H-pyrrolo[1,2-b]pyrazol-4-one, Molport-022-469-240. Parting the nitrogens is energetically favorable: for pyrazole H_f_(s) = +105 kJ/mol,

H_f_(g) = +179 kJ/mol; for imidazole H_f_(s) = +49 kJ/mol, H_f_(g) = +132.9 kJ/mol [Haynes 2013]. Moreover, the original His has an imidazole and not a pyrazole ring, and so does the residue.

## 5 Discussion

### Entropy of decomposition

In the tables ofDomalski [1972],Chickos and Acree [2002] and Acree and Chickos [2010], at temperatures coinciding with our peak temperatures, “Heats of sublimation” of the order of our endothermic peak areas are reported. Our QMS signals prove that chemical decomposition is involved, but that should have been obvious from the DSC data alone. The average of the entropies of transformation, S_peak_ = H_peak_/T_peak_, is 215 J/Kmol, way above the usual entropies of melting (22 J/kmol for H_2_O, 28 for NaCl, 36 for C_6_H_6_), and higher than typical entropies of evaporation (41 J/Kmol for H_2_O, 29 for CS_2_, 23 for CO_2_, 21 for NH_3_). The endothermic heats in the peaks are therefore neither enthalpies of fusion nor enthalpies of sublimation, they are heats of reaction accompanied by phase changes. There is transformation and decomposition, but no reversible melting. Amino acids are stable in solid form, but not as liquids or gases.

### Peptide bond formation

Five of the eight amino acids have residues containing peptide bonds, –C(=O)-NH–, only Asp and Asn leave polysuccinimide (PSI), Cys leaves cyclic pyrrolines. The preponderance of water in thermal decomposition is not surprising. In natural protein formation, each participating amino acid suffers damage. In the condensation reaction, where the N-terminus of one molecule reacts with the C-terminus of its neighbour, the planar peptide bond –Cα-CO-N-Cα– is formed. The N-atoms on, and the keto-bound O-atoms off the backbones retain their position. H_2_O is ejected, but neither NH_3_ nor CO_2_ are produced in protein formation. Thermal decomposition of amino acids is analogous. In protein formation, the endothermic heat is provided by ATP, in amino acid decomposition it is thermal energy.

**Figure 5.**
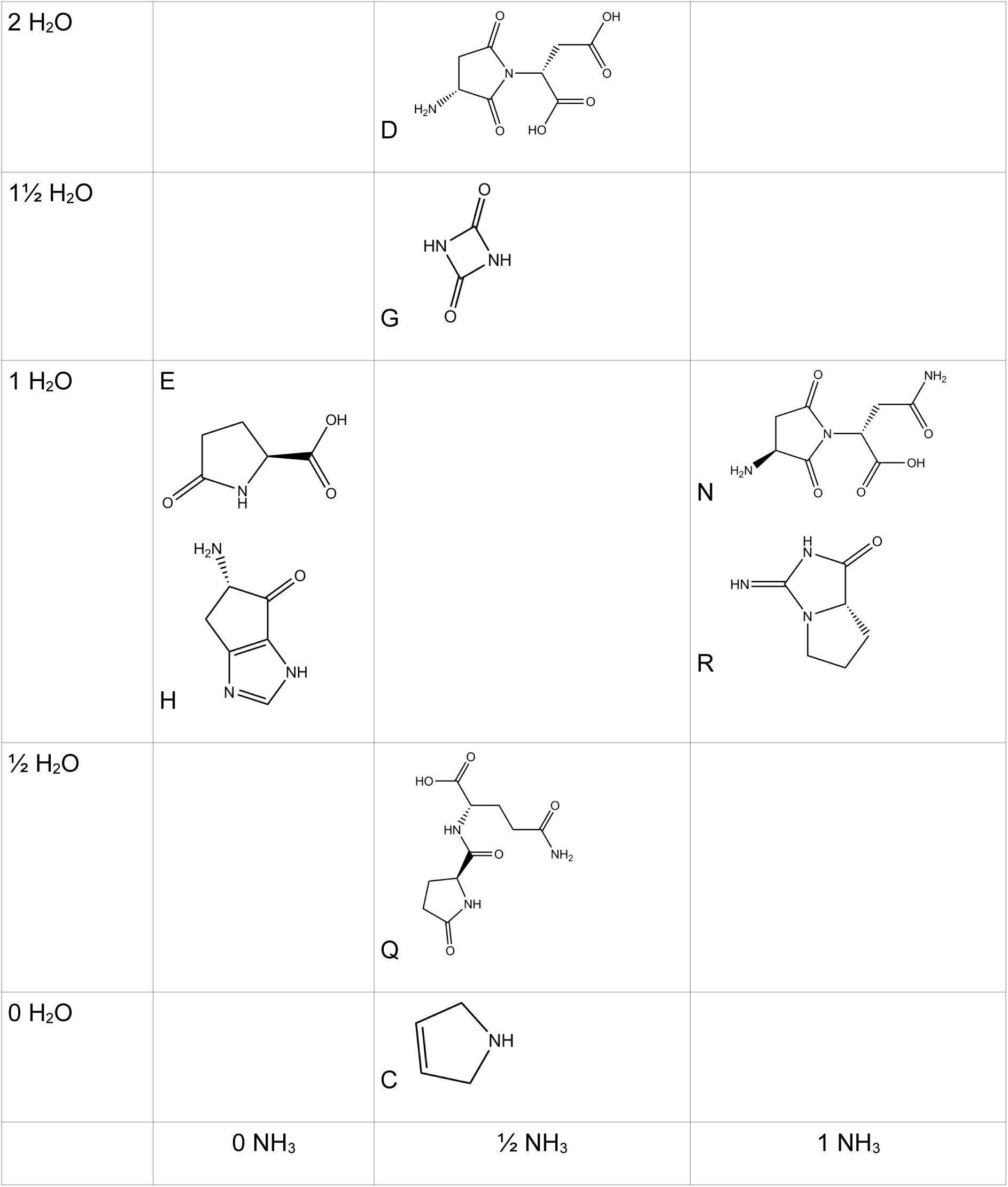
All residues are obtained by ejection of 0, V, 1, 1 % or 2 mols of H_2_O or NH_3_, placing them on two axes. All residues contain either 0, V or 1 mol NH_3_ or H_2_O, placing them on two axes. Most of them contain peptide bonds. The polysuccinimide of D and N is an exception, cysteine, for lack of oxygen, the other.

### Peak areas

Quantitatively, the parallel between protein formation and pyrolysis is confirmed on the enthalpy level. In the formation of a dipeptide, X+Y ⇾ H_2_O + (X/Y), the difference between the enthalpies of the reactants and the products go into the formation of the peptide bond: H_f_(X) + H_f_(Y) = −242 kJ + H_f_(X/Y) + H(PB). With the tabulated value [Haynes 2013,Domalski 1972] for H_f_(X), H_f_(Y) and H_f_(X/Y) one calculates H(PB) = −67 kJ in glycylglycine, −70 kJ in alanylglycine, −43 kJ in serylserin, −78 kJ in glycylvaline, −65 kJ in leucylglycine, −91/2 kJ in triglycylglycine, −86/2 kJ in leucylglycylglycine, and −58 kJ in glycylphenylalanine. The average value is −59±13 kJ per peptide bond. The narrow standard deviation indicates that the enthalpy of forming a peptide bond is insensitive to its environment, therefore the endothermic values of oligomerization or polymerization should be close to this.

One concludes that the formation of a peptide bond in a linear dimer is endothermic with an enthalpy of 59±13 kJ. It is tempting to compare this with the areas of the DSC peak, the observed endothermic heat of the decomposition reaction. The average of the eight amino acids is −105±27 kJ/mol. One concludes that essentially the endothermic heat of decomposition, the peak area, goes into peptide bond formation.

### NH3 Production

In cases where the N-terminus, untouched by the condensation, remains attached to a cyclic product, it could be cut off as NH_3_, contributing up to % mol NH_3_/mol AA.

Remarkable is the absence of methane (CH_4_, 16 Da), hydrogen cyanide (HCN, 27 Da) and formamide (CH_3_NO, 45 Da), all in mass channels where we would have seen them. These, suspected in prebiotic synthesis of amino acids, do not appear in their decomposition.

Although at most only three molecules are involved, two gases and one monomolecular residue, identification of the structure of the latter is not unequivocal, there remain more or less probable other possibilities than our choices. Clearly, without QMS, data from DSC and TGA could not possibly suffice to identify decomposition chains.

## 6 Conclusion

Amino acids decompose thermally, they do not sublimate, nor do they melt. Only three gases are formed, mostly H_2_O, less so NH_3_ and hardly any CO_2_. In all amino acids investigated, Gly, Cys, Asn, Asp, Gln, Glu, Arg, His, the liquid or solid residues are lactams and heterocyclic compounds with 5 or 6-membered non-(or only partially) aromatic rings, containing one or two nitrogen atoms (pyrrolidines, piperidines, pyrrazolidines, piperazines), most of them with peptide bonds present.

## Acknowledgements

We thank Angela Rutz and Frederik Schweiger for technical assistance. This work would have been impossible without the continuous support of Eduard Arzt.

## References

Acree W, Chickos JS (2010) Phase Transition Enthalpy Measurements of Organic and Organometallic Compounds. Sublimation, Vaporization and Fusion Enthalpies From 1880 to 2010. J Phys Chem Ref Data 39

Acree W, Chickos JS (2016) Phase Transition Enthalpy Measurements of Organic and Organometallic Compounds. Sublimation, Vaporization and Fusion Enthalpies From 1880 to 2015. Part 1. C-1-C-10. J Phys Chem Ref Data 45

Banack SA, Metcalf JS, Jiang LY, Craighead D, Ilag LL, Cox PA (2012) Cyanobacteria Produce N-(2-Aminoethyl)Glycine, a Backbone for Peptide Nucleic Acids Which May Have Been the First Genetic Molecules for Life on Earth. Plos One 7

Chickos JS, Acree WE (2002) Enthalpies of sublimation of organic and organometallic compounds. 1910-2001. J Phys Chem Ref Data 31: 537–698

Domalski ES (1972) Selected Values of Heats of Combustion and Heats of Formation of Organic Compounds Containing the Elements C, H, N, O, P, and S. J Phys Chem Ref Data 1: 221–277

Follmann H, Brownson C (2009) Darwin’s warm little pond revisited: from molecules to the origin of life. Naturwissenschaften 96: 1265–1292

Haitinger L (1882) Vorläufige Mittheilung über Glutaminsaäure und Pyrrol. Monatshefte für Chemie und verwandte Teile anderer Wissenschaften 3: 228–229

Haynes WM (2013) Section 5: Standard Thermodynamic Properties of Chemical Substances. In CRC Handbook of Chemistry and Physics, Boca Raton, FL, USA: CRC Press, Taylor and Francis Group

Kim et al (2017) Electron-Impact Cross Section Database; http://phvsics.nist.gov/PhvsRefData/ASD/ionEnergy.html. In Gaithersburg, MD, USA: NIST

Mosqueira FG, Ramos-Bernal S, Negron-Mendoza A (2008) Prebiotic thermal polymerization of crystals of amino acids via the diketopiperazine reaction. Biosystems 91: 195–200

Rodante F, Marrosu G, Catalani G (1992) Thermal-Analysis of Some Alpha-Amino-Acids with Similar Structures. Thermochim Acta 194: 197–213

Schiff H (1897) Ueber Polyaspartsäuren. Berichte der deutschen chemischen Gesellschaft 30: 2449–2459

